# Socio-spatial heterogeneity in participation in mass dog rabies vaccination campaigns, Arequipa, Peru

**DOI:** 10.1101/542878

**Authors:** Ricardo Castillo-Neyra, Amparo M. Toledo, Claudia Arevalo-Nieto, Hannelore MacDonald, Micaela De la Puente-León, Cesar Naquira-Velarde, Valerie A. Paz-Soldan, Alison M. Buttenheim, Michael Z. Levy

**Affiliations:** Department of Biostatistics, Epidemiology and Informatics, Perelman School of Medicine, University of Pennsylvania Perelman School of Medicine, Philadelphia, Pennsylvania, USA; Zoonotic Disease Research Lab, One Health Unit, School of Public Health and Administration, Universidad Peruana Cayetano Heredia, Arequipa, Peru; Department of Biology, School of Arts and Sciences, University of Pennsylvania, Philadelphia, Pennsylvania, USA; Global Community Health and Behavioral Sciences, Tulane School of Public Health and Tropical Medicine, New Orleans, Louisiana, USA; Department of Family and Community Health, School of Nursing, University of Pennsylvania, Philadelphia, Pennsylvania, USA

**Keywords:** Community participation, Disease outbreaks, Dog diseases, Immunization programs, Mass vaccination, One Health, Rabies, Vaccination coverage, Zoonoses

## Abstract

To control and prevent rabies in Latin America, mass dog vaccination campaigns (MDVC) are implemented mainly through fixed-location vaccination points: owners have to bring their dogs to the vaccination points where they receive the vaccination free of charge. Dog rabies is still endemic in some Latin-American countries and high overall dog vaccination coverage and coverage evenness are desired attributes of MDVC to halt rabies virus transmission. In Arequipa, Peru, we conducted a door-to-door post-campaign survey on >6,000 houses to assess the placement of vaccination points on these two attributes. We found that the odds of participating in the campaign decreased by 16% for every 100 m from the owner’s house to the nearest vaccination point (p=0.041) after controlling for potential covariates. We found social determinants associated with participating in the MDVC: for each child under 5 in the household, the odds of participating in the MDVC decreased by 13% (p=0.032), and for each decade less lived in the area, the odds of participating in the MDVC decreased by 8% (p<0.001), after controlling for distance and other covariates. We also found significant spatial clustering of unvaccinated dogs over 500 m from the vaccination points, which created pockets of unvaccinated dogs that may sustain rabies virus transmission.

Understanding the barriers to dog owners’ participation in community-based dog-vaccination programs will be crucial to implementing effective zoonotic disease preventive activities. Spatial and social elements of urbanization play an important role in coverage of MDVCs and should be considered during their planning and evaluation.

**Author summary:** In Peru and other dog rabies-affected countries, mass dog vaccination campaigns (MDVC) are implemented primarily through fixed-location vaccination points: owners have to bring their dogs to the vaccination points where they receive the vaccination. To stop rabies virus transmission, a high and even dog vaccination coverage is desired. In Arequipa, Peru, following a MDVC, we conducted a door-to-door survey of >6,000 houses to assess how the placement of vaccination points affected coverage of the campaign. When comparing dog owners with similar characteristics, we found that the odds of participating in the MDVC was reduced by 16% for every 100 m distance from the nearest vaccination point. Some social conditions were also associated with participating in the MDVC: for each child under 5 in the household, odds of participating in the MDVC decreased by 13%, and for each decade less lived in the area, the odds of participating in the MDVC decreased by 8%. Distance to the vaccination point and variation in social conditions across the city play important roles in achieving coverage of MDVCs and should be considered during campaign planning and evaluation.

## 1. Introduction

The city of Arequipa is in the midst of a sustained dog rabies outbreak. The introduction of rabies virus into the city has been ascribed to the unintentional transport of rabid dogs from the rabies-endemic state of Puno during human migration [1–3], and the persistence of transmission is likely due to low coverage in the annual city-wide dog vaccination campaigns [3]. Following the detection of the outbreak in Arequipa city in 2015, the Ministry of Health of Peru (MOH) initiated additional vaccination campaigns in the city with varying intensity [4]. These additional efforts have not quelled the epidemic: more than 160 rabid dogs have been detected as of 2019.

Epidemics of dog rabies are ongoing in major urban centers across Latin America and worldwide [1,5–8]. Since bites from rabies-infected dogs cause 99% of human rabies deaths [9], the control and elimination of dog-mediated human rabies relies on a population-wide animal-centered strategy: mass dog rabies vaccination [8,10,11]. Dog vaccination has dramatically decreased the global burden of human rabies since 1955 [5,12,13]; in the Americas, national programs centered around mass dog vaccination have achieved enormous advances [8,14,15], reducing the incidence of dog rabies by 98% since 1983 [14]. In most rabies-affected countries, government health entities (e.g. MOH) organize annual MDVC that are held in outdoor settings. These campaigns are usually free of charge and voluntary [8,16] and campaign promotion varies greatly in format, content and intensity [1,17,18].

There are three non-mutually exclusive strategies used in Peru to implement MDVC: fixed vaccination posts, mobile teams setting up a temporary mobile post or conducting ‘street vaccination’, and door-to-door vaccination [19]. For the fixed-point strategy, the vaccinators wait for the dog owners to bring the dogs to a unique place. For the mobile team strategy, the vaccinators move from one location to another during the day, vaccinating dogs along their way and spending short periods (i.e. under an hour) in a location before moving on, waiting for the dog owners to bring the dogs to these moving locations. For the door-to-door strategy, vaccinators knock on doors asking to vaccinate dogs in the household. Locations of the fixed vaccination sites are typically determined by a combination of convenience and prominence of the location (e.g. the entrance to a health post, a well-known park) [11]. In Peru, routes for door-to-door and mobile team approaches may or may not be decided in advance, and teams may move during the course of the day looking for dogs along their routes.

The fixed-point strategy has been extensively used in Latin America and Africa, even though it has frequently failed to attain coverage targets [1,8,20–24]. The main reasons for its extensive application are that fixed-point vaccination is easier to implement and less costly than other strategies [18]. In many cases fixed-point is combined with other strategies, particularly when initial activities are unsuccessful [19,23,25]. However, high dog owner participation in MDVC and other dog-centered health campaigns (e.g. de-worming dogs to prevent human hydatid disease) has proven difficult to achieve in many areas [24,25]. It is necessary to understand barriers to community-based control strategies targeting dog populations in dog rabies-affected countries where coverage does not reach the minimum 70% recommended by the World Health Organization to attain herd immunity [11], much less the 80% recommended by the Pan-American Health Organization for the region [26].

In cities, the social and spatial aspects of urbanization can facilitate the emergence of dog rabies and complicate its control [27,28]. In Arequipa, the locations of rabid dogs have been associated with urban structures [27], and dog owners from areas with different levels of urbanization have reported distinct correlates of vaccinating their dogs against rabies [1]. The changing urban landscape and social processes in rapidly-growing cities have been associated with uptake of health-related services [29–32] and may be related to the low dog vaccination coverage in Arequipa. The objectives of the present study were to quantitatively assess the association between distance to a vaccination point and dog owner participation in mass dog vaccination campaigns in an urban setting, and to evaluate the effect of such distance on overall vaccination coverage and evenness.

## 2. Methods

### 2.1. Ethics statement

Ethical approval was obtained from Universidad Peruana Cayetano Heredia (approval number: 65369), Tulane University (approval number: 14–606720), and University of Pennsylvania (approval number: 823736). All human subjects in this study were adults.

### 2.2. Study setting

The study was conducted in Alto Selva Alegre (ASA) (human population for 2015: 82,412; density: 11,902 people/km^2^), one of the 14 districts of the city of Arequipa. Arequipa, Peru’s second largest city, is home to 969,000 people and is situated at ~2,300 meters above sea level. The first detection of a rabid dog in the city of Arequipa occurred in March 2015 in ASA. By June 2016, when our data were collected, 43 rabid dogs had been detected in 8 districts, but it is assumed that number represents a small fraction of the total number of cases [27]. The city of Arequipa comprises communities spanning different stages of urbanization and different migration histories, from old established neighborhoods, to young neighborhoods, to recent invasions [33]. Within this gradient of development, young neighborhoods and recent invasions are often located on the periphery of the city (peri-urban area) and the older localities are nearer to the center (urban area) [33]. Compared to the urban area, peri-urban areas generally have lower socioeconomic status, fewer community resources, more security problems, and often more rugged and uneven terrain (Figure 1). As new neighborhoods mature into established neighborhoods with wealthier residents, homes are improved with better quality construction material and permanent utility connections, and connectivity with the rest of the city increases with better sidewalks, roads, and transportation access. ASA transects the city, running from the center to the periphery, and the district continues to grow towards the outskirts of the city. In our study, participants represented either the urban or peri-urban areas of the city of Arequipa. We included 21 urban neighborhoods founded many decades ago, and 9 peri-urban neighborhoods that originated around 2000 or later.

**Figure 1.**
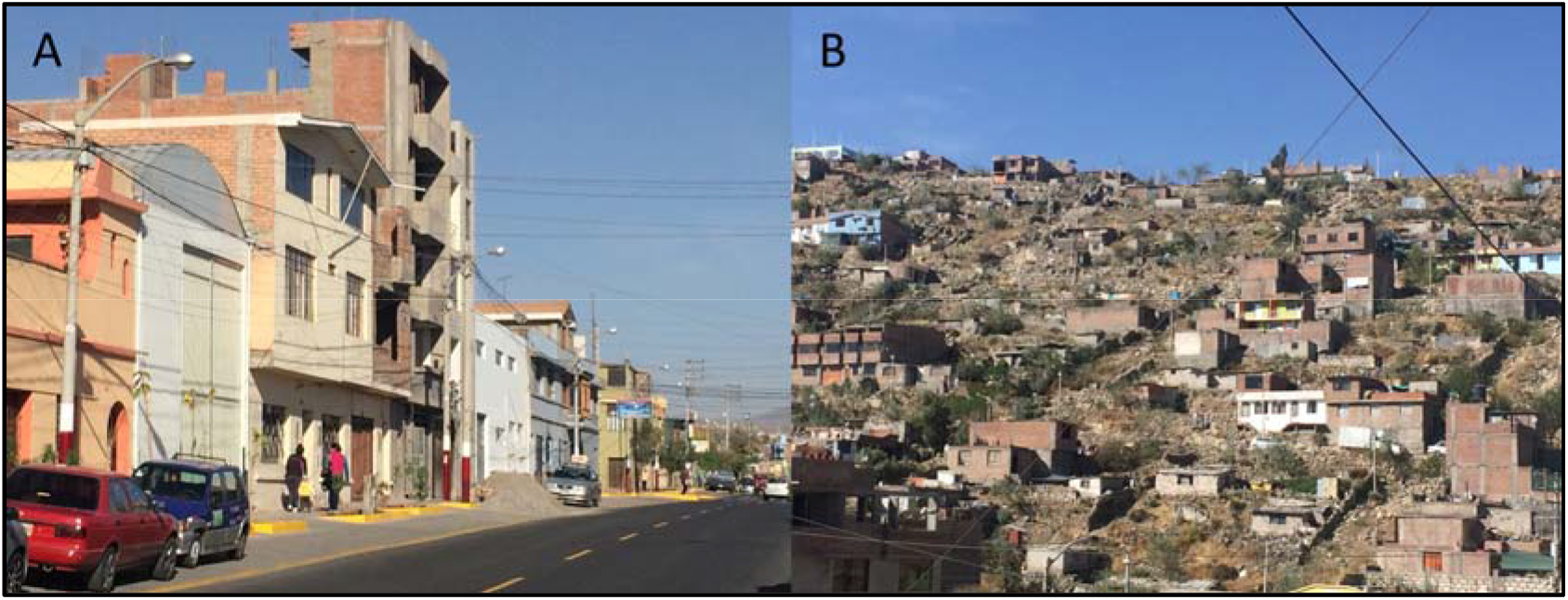
Study communities display landscape heterogeneity. A: Urban area. B: Peri-urban.

In ASA, the MOH conducted a mass dog vaccination campaign in June 2016. A detailed description of the mass dog vaccination campaigns can be found elsewhere [1], and the dog population estimator used in Arequipa can be found in [27].

### 2.3. Data collection

In collaboration with the MOH, we georeferenced every stationary and mobile vaccination team during the three weekends when the campaign was implemented in ASA. Due to volunteer assistance, some dogs were vaccinated in weekdays during the campaign; we did not georeference the location of those volunteering vaccinators, mainly because their schedule was haphazard and unpredictable. We started door-to-door surveys immediately after the vaccination campaign, visiting every household in the study area and consenting and surveying only one adult (≥18) per household.

The door-to-door survey was designed based on the rabies literature and based on our qualitative studies of local communities, in which we found specific household-level barriers to vaccination [1]. We collected household variables (e.g. number of household members; number of children under 5 years old), dog owner or interviewee variables (e.g. gender; educational attainment), and dog variables (e.g. vaccination status; age). All houses in the study localities were geocoded and the survey data were linked to the household coordinates. We estimated the Euclidean distance between households and the closest vaccination point (fixed, mobile, or either).

### 2.4. Statistical Analysis

We estimated the total vaccination coverage in the study area and compared the coverage in urban vs. peri-urban localities with a chi-squared test. We estimated the human to dog ratio and bootstrapped it 10,000 times to estimate its confidence interval. To evaluate the baseline characteristics of households and dog owners by participation in the MDVC, we defined an ordinal outcome for houses with dogs: *no participation* (no dog vaccinated in the house), *partial participation* (some, but not all dogs in the house, vaccinated), and *full participation* in the MDVC (all dogs in the house vaccinated). We used a chi-square test to compare categorical variables with 10 or more observations per group, Fisher’s exact test for categorical variables with fewer than 10 observations in any subgroup, and Mann–Whitney U test for age of the dog owner or interviewee, which did not follow a normal distribution and was truncated at 18 years. We compared the individual characteristics of vaccinated and unvaccinated dogs with chi-square for categorical variables and with Mann–Whitney U test for dog’s age.

Our main objective was to assess the association between distance to the vaccination point and participation in the MDVC. For distance to the vaccination point, we used the Euclidean distance from the dog’s house to the closest vaccination point, either fixed or mobile. For participation in the MDVC we used the ordinal values described above: *no participation*, *partial participation*, and *full participation.* We compared proportional odds logistic regression (POLR), non-proportional odds logistic regression, and multinomial regression. The ordinal models were superior to the multinomial regression, and the proportional odds assumption holds for most of the covariates. Given that the categories of participation are inherently ordinal and that providing a single point estimate per covariate is more interpretable, the POLR model was favored. Based on the recent literature and our local studies [1,34,35], the following covariates were used for model building: having a dog leash at home, number of children under 5 years old at home, time living in the area, rabies status of the last place they lived in before living in the study area, number of dogs at the house, age and gender of the dog owner, and educational attainment. We considered transformed distance to capture non-linear effects and interactions between distance and having a leash. The fit of the alternative models to the data was compared with Akaike’s Information Criteria (AIC). We also attempted to build a hierarchical model to take into account the spatial autocorrelation within locality. However, given that within each locality there was at most one vaccination point, the variable distance from the house to the vaccination point would be unidentifiable under such hierarchical model. The final POLR model fitted with the R package MASS [36] was:

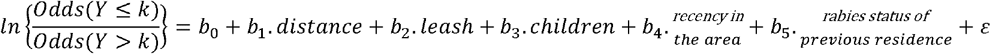

where *k* takes the values 0 (no participation), 1 (partial participation), and 2 (full participation). All statistical tests were 2-sided, and significance level was 0.05.

We tested the spatial pattern of vaccinated and unvaccinated dogs in relation to vaccination tents for clustering using the bivariate cross K-function. This function estimates spatial dependence between two types of points (i.e. unvaccinated dogs and vaccination points) by measuring the expected number of points of type *i* within a given distance to a point of type *j* divided by the overall density of the points of type *i*. We used the Kcross function in the R package spatstat [37] to estimate deviations between the K function estimated for our data and the theoretical K function corresponding to a completely random Poisson point process for vaccinated dogs to tents and unvaccinated dogs to tents. Deviations greater than the theoretical K function indicate that the mean point count is higher than expected under complete spatial randomness (CSR) and thus some degree of clustering is present between the two event types at the indicated distance. Similarly, deviations less than the theoretical K function indicate that the mean point count is lower than expected under CSR which therefore indicates that some degree of dispersion is present between the two event types at the indicated distance.

In order to investigate the association between geolocation and the odds of canine vaccination we fitted Generalized Additive Models (GAMs) to our data using the R package MapGAM [38]. GAMs are an extension of linear regression models in which both parametric and non-parametric terms are used to estimate the outcome of interest. We used a two-dimensional locally weighted smooth (LOESS) of latitude and longitude for our non-parametric term. The LOESS smoother fits each data point by weighting it towards nearby points, where weighting is based on the distance to the point being fitted. The percentage of data points in the region that will be used to predict a particular point is referred to as the span. The optimal span size used for smoothing was determined by minimizing the AIC. We mapped the odds ratio for each point on pre-specified grids of each locality (from polygon data) and next tested the null hypothesis that the odds of each points’ vaccination status did not depend on geolocation using permutation tests. For each test, the paired latitude and longitude coordinates were randomly permuted but vaccination status was held fixed. 1000 permutations were run for each locality and contour lines encircle areas with significantly increased or decreased vaccination odds as indicated by point wise p-values computed from the permutation ranks. All models and figures were created with R [39].

## 3. Results

Based on our survey, the estimated vaccination coverage of the MOH MDVC was only 58.1%, and it was low in both urban and peri-urban localities (58.0% vs. 58.6% respectively, chi2=0.086, p=0.769). Only 3.4% of dogs were (reportedly) vaccinated in private clinics, bringing our estimated total coverage to 61.5%. Participation in our survey was higher in the urban area (88.8%) compared to the peri-urban area (61.6%) (mean= 82.0%, chi2=6458.5, p<0.001). The total number of dogs in the surveyed houses was 5,292 and the human to dog ratio was 3.78:1 (95% CI: 3.69:1 – 3.89:1). In total, 65.3% of surveyed houses had dogs, but this number was higher in peri-urban areas (70.0% compared to 64.6% in urban areas, chi2=6.529, p=0.011). For 76.9% of vaccinated dogs in the area, the person who took them to the MDVC is the person who responded to the survey. In our study area the urbanization process involves new localities being founded and settlers moving in. Accordingly, we found that the time of residency (or the year people moved into this area) was clustered at the locality level. However, we found that people living in or founding a locality do not necessarily share the same place of origin or previous residence.

When we compared the distance from each house to the closest fixed vaccination point, to the closest mobile vaccination point, and to either the closest fixed or mobile vaccination point, we found a clear gradient in distance from vaccination point from non-participant houses (farther) to houses that partially participated (closer) to houses that fully participated (closest) (Table 1). The proportion of households with children under 5 was higher in households that did not participate in the campaign (36%) compared to houses that participated fully or partially in the campaign (31% and 29% respectively). The proportion of houses with a dog leash increases from those who did not participate, to those who participated partially, to those that fully participated of the MDVC. There were some differences in MDVC participation by migration history: people who have lived longer in ASA tend to report higher participation in the MDVC compared to those who have lived fewer years in ASA. Also, there is a slight difference in participation in the MDVC depending on rabies status of previous residence, with more people participating in the campaign, partially or fully, if they were migrants from a rabies endemic area (Table 1). Other variables, such as educational attainment, the proportion of female dog owners or interviewees, and the proportion of households in urban localities were all similar in those households that participated fully or partially in the campaign compared to those households that did not participate of the campaign (Table 1). Having multiple dogs is a prerequisite to be in the partial participation group; therefore, houses with partial participation had on average more dogs, but the number of dogs per house was very similar in houses that participated fully compared to those that did not participate in the campaign (Table 1). Those who did not participate in the MDVC reported more frequently not knowing about the campaign before it happened (Table 1), but many of them reported learning about the campaign the same day it occurred. In S1_Table, we report the media channels through which they learned about the campaign either after it occurred or the same day it occurred.

**Table 1.** Characteristics of the dog owner population in the study area by participation in the MDVC, Arequipa City, Peru, 2016.

| Interviewee/Owner Variables | Vaccinated no dog (n=1,017) | Vaccinated some dogs (n=176) | Vaccinated all dogs (n=1,434) | p |
| --- | --- | --- | --- | --- |
| Distance to any closest vaccination point – Mean (SD) | 125m (96) | 117m (114) | 103m (83) | <0.001<sup>a</sup> |
| Distance to closest fixed vaccination point – Mean (SD) | 501m (479) | 490m (420) | 461m (422) | 0.119<sup>a</sup> |
| Distance to closest mobile vaccination point – Mean (SD) | 137m (101) | 127m (119) | 114m (93) | <0.001<sup>a</sup> |
| Number of dogs in houses with dogs – Mean (SD) | 1.79 (1.10) | 2.96 (1.24) | 1.80 (1.05) | <0.001<sup>a</sup> |
| Houses with children under 5 | 36.4% | 28.9% | 30.8% | 0.019<sup>b</sup> |
| Age of dog owner/interviewee – Median (IQR) | 39 years (28 - 52) | 40 years (29 - 50) | 42 years (30 - 53) | 0.024<sup>c</sup> |
| Time living in ASA if migrant – Mean (SD) | 18.1 years (13.7) | 21.2 years (13.9) | 22.1 years (14.8) | <0.001<sup>a</sup> |
| Area before living in ASA |  |  |  | 0.036<sup>b</sup> |
| Rabies epidemic (Arequipa) | 90.7% | 94.1% | 89.8% |  |
| Rabies endemic (Puno) | 1.6% | 2.4% | 3.3% |  |
| Rabies free (rest of country) | 7.7% | 3.6% | 6.9% |  |
| Educational attainment |  |  |  | 0.509<sup>b</sup> |
| Illiterate | 0.6% | 0.5% | 0.6% |  |
| Primary school | 11.5% | 11.4% | 13.6% |  |
| High school | 43.2% | 48.3% | 41.1% |  |
| Technical academy | 21.2% | 15.3% | 20.6% |  |
| University | 22.3% | 23.3% | 23.3% |  |
| Preferred not to respond | 1.2% | 1.1% | 0.7% |  |
| Owens at least one dog leash | 51.0% | 54.5% | 60.1% | <0.001<sup>b</sup> |
| Female interviewees/owners | 64.6% | 73.3% | 64.4% | 0.032<sup>b</sup> |
| Did not know about the campaign before it happened | 15.4% | 2.3% | 1.3% | <0.001<sup>b</sup> |
| Live in urban locality (vs. peri-urban) | 78.8% | 78.1% | 79.4% | 0.002<sup>b</sup> |

*p* values estimated with <sup>a</sup> one-way ANOVA; <sup>b</sup> Chi square test; and <sup>c</sup> Kruskal Wallis test.

Compared to vaccinated dogs, unvaccinated dogs were older, were more likely female, had more free access to the street, had owners with no leash for them at home, and were less likely to be walked. Multipurpose dogs (dogs reported as guard and company dogs) were more likely to be vaccinated (Table 2). Based on the stated source of the dog, those received as gifts are more likely to be vaccinated and dogs born at home or adopted/picked on the street are less likely to be vaccinated (Table 2). Being considered purebred or being spayed/neutered was not associated with dog vaccination status (Table 2).

**Table 2.** Dog characteristics by vaccination status in the MDVC (dogs vaccinated privately not included).

| Dog Variables | Unvaccinated dog<br>(n=2,019) | Vaccinated dog<br>(n=2,800) | p |
| --- | --- | --- | --- |
| Age of dogs – Mean (SD) | 24 months (12 – 60) | 20 months (10 – 54) | 0.039<sup>a</sup> |
| Female dogs | 42.1% | 36.8% | <0.001<sup>b</sup> |
| Free access to the street | 20.6% | 25.7% | 0.002<sup>b</sup> |
| Leash for the dog | 33.8% | 45.3% | <0.001<sup>b</sup> |
| Dogs that are walked, either with or without leash. | 55.9% | 66.9% | <0.001<sup>b</sup> |
| Dog function |  |  | <0.001<sup>b</sup> |
| Company | 52.0% | 49.0% |  |
| Protection | 23.4% | 20.9% |  |
| Company & Protection | 24.6% | 29.6% |  |
| Breeder/business | 0.0% | 0.5% |  |
| Dog source |  |  | <0.001<sup>b</sup> |
| Gift | 51.2% | 56.5% |  |
| Bought at market/store | 17.4% | 18.3% |  |
| Born at home | 15.6% | 12.3% |  |
| Adopted/Picked on the street | 12.3% | 10.8% |  |
| Bought from friend/neighbor | 2.6% | 2.0% |  |
| Do not know | 1.0% | 0.0% |  |
| Purebred as reported by interviewee | 25.7% | 24.9% | 0.589<sup>b</sup> |
| Spayed/Neutered | 3.5% | 2.6% | 0.157<sup>b</sup> |

*p* values estimated with <sup>a</sup> Student's t-test; and <sup>b</sup> Chi square test.

In the multivariable regression analysis, we found that distance to the vaccination site is strongly associated with participation of the MDVC. The odds of not participating in the MDVC were 16% lower for someone who lived 100 meters farther from the vaccination point after adjusting for other covariates and this difference was statistically significant (Table 3). Those having a dog leash at home had 35% higher odds of participation of the MDVC, either fully or partially, compared to those who did not have a dog leash after adjusting for other covariates. The odds of participating in the MDVC, either fully or partially, were 13% lower for each additional child under 5 years at home, after adjusting for other covariates. Migration history was associated with participating in the MDVC; participation was lower in those who migrated more recently to the study area (8% lower odds of participation for each decade less lived in the area, after adjusting for other covariates). Another component of migration history, the previous residence region, was also associated with participating in the MDVC: those whose previous residence was a rabies-free region or was any district within Arequipa were 23% to 32% less likely to participate of the campaign compared to those whose previous residence was a rabies-endemic region, after adjusting for other covariates (Table 3). Demographic variables such as owner’s/interviewee’s age, gender and educational attainment, and other household-level variables, such as number of dogs at the house, were dropped during model selection because they neither improved the fit of the model nor were statistically associated with participating in the MDVC.

**Table 3.**
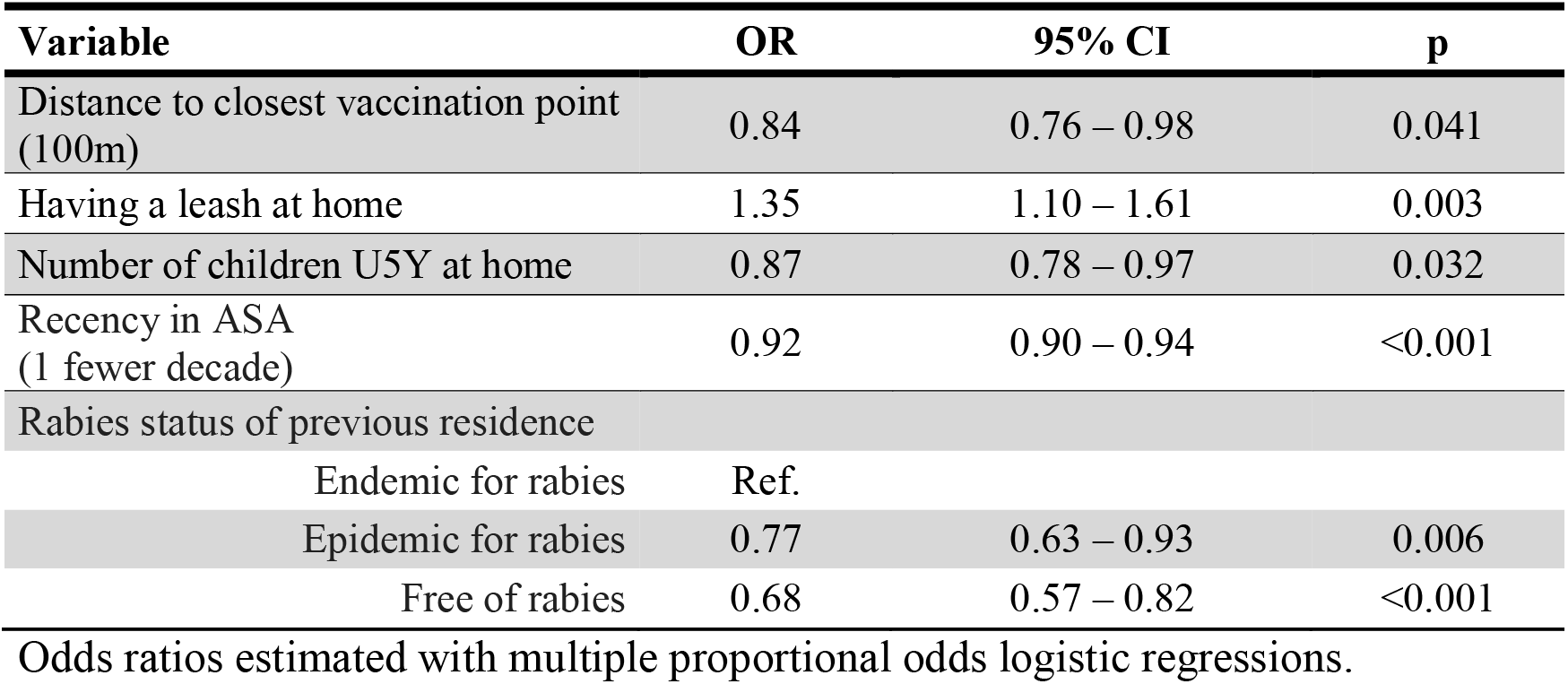
Factors associated with participating in the Mass Dog Vaccination Campaign, Arequipa City, Peru, 2016.

| Variable | OR | 95% CI | p |
| --- | --- | --- | --- |
| Distance to closest vaccination point (100m) | 0.84 | 0.76 – 0.98 | 0.041 |
| Having a leash at home | 1.35 | 1.10 – 1.61 | 0.003 |
| Number of children U5Y at home | 0.87 | 0.78 – 0.97 | 0.032 |
| Recency in ASA (1 fewer decade) | 0.92 | 0.90 – 0.94 | <0.001 |
| Rabies status of previous residence |  |  |  |
| Endemic for rabies | Ref. |  |  |
| Epidemic for rabies | 0.77 | 0.63 – 0.93 | 0.006 |
| Free of rabies | 0.68 | 0.57 – 0.82 | <0.001 |

Odds ratios estimated with multiple proportional odds logistic regressions.

The significant association between distance to the vaccination point and odds of participating in the MDVC has consequences for the distribution of unvaccinated dogs in the area. We observed spatial clustering of unvaccinated owned dogs as a function of the distance from the house to the vaccination point. These pockets of unvaccinated dogs closer to each other than expected by chance occur at 500 meters from the vaccination point or further (Figure 2). We also analyzed the spatial odds of participating in the MDVC, that is, the association between their specific geolocation and the vaccination point. For areas served by fixed-point vaccination, there was a clear smooth spatial effect with higher odds of participating for houses closer to the vaccination point and a decreasing gradient farther away from the vaccination point. The spatial effect of the fixed-point vaccination strategy creates two clearly defined zones: a large zone with statistically significantly high odds of participation in the MDVC and another large zone with statistically significantly low odds of participation in the MDVC (Figure 3). For areas served by mobile teams, there were more spots of significant low and high odds of participation in the MDVC and these spots were spread in the study area without a clear association between the spots and the locations where the vaccination teams stopped to wait for dogs or to vaccinate dogs (Figure 3).

**Figure 2.**
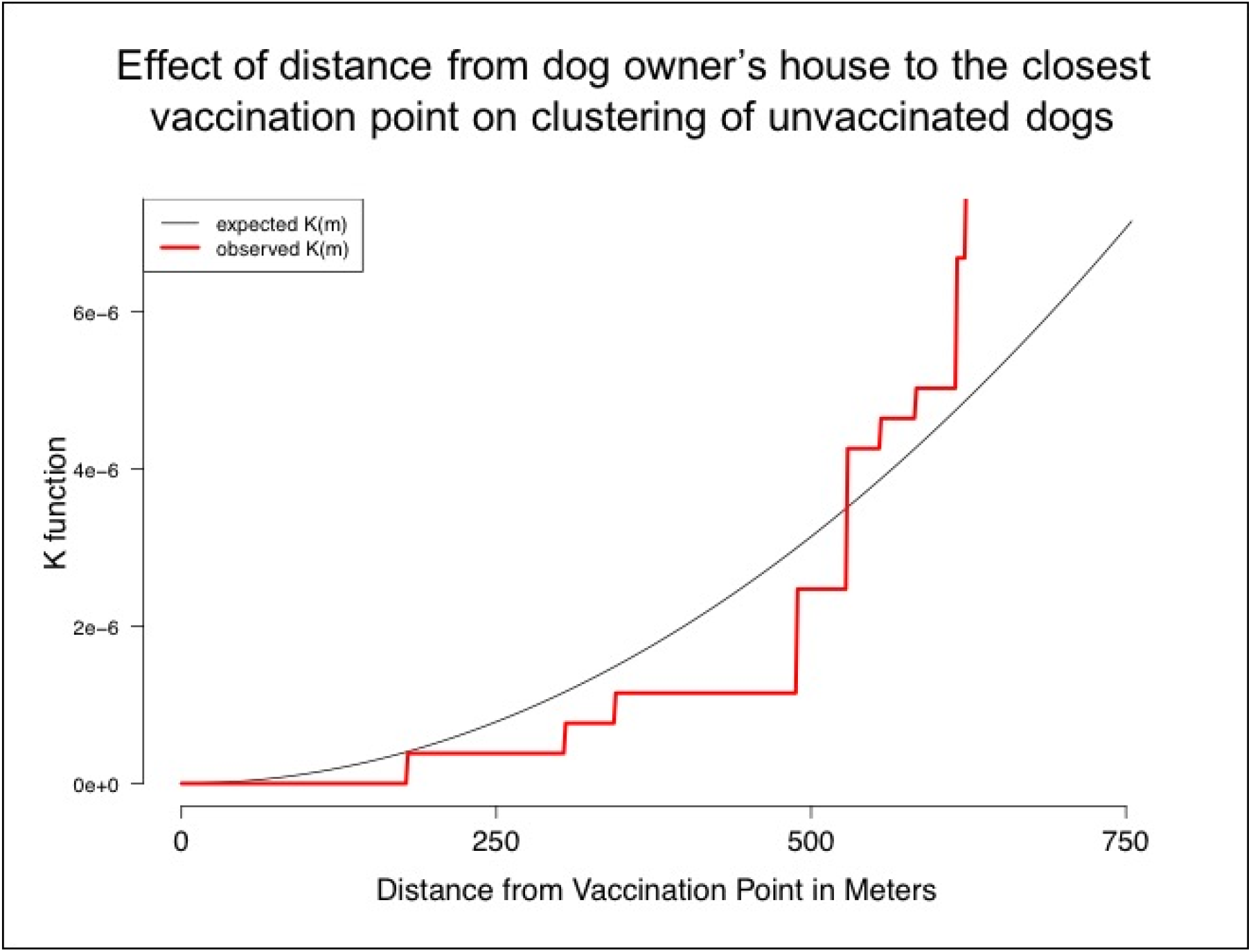
Clustering of unvaccinated dogs as a function of distance from the vaccination point.

**Figure 3.**
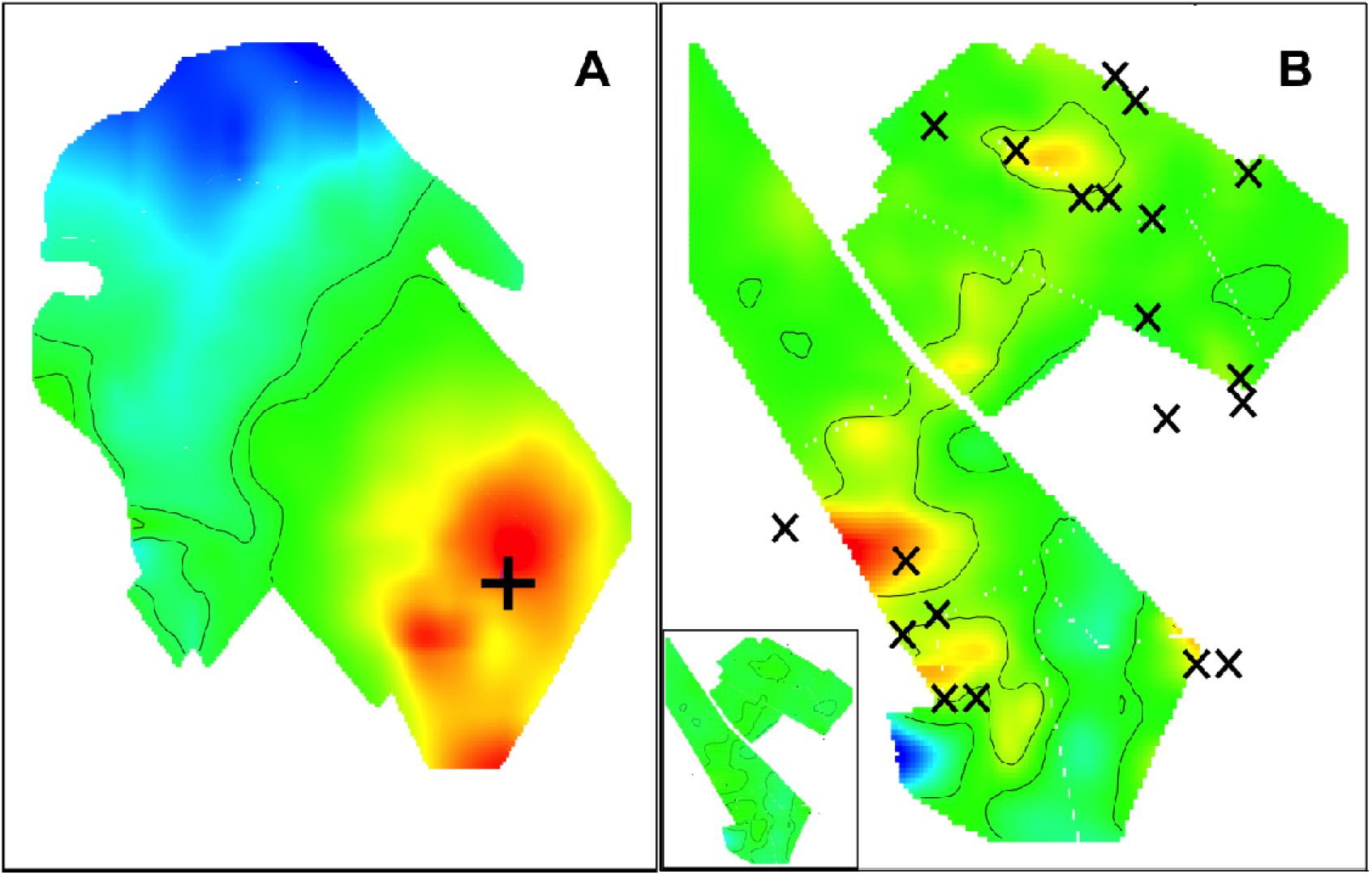
Spatial odds ratios for participating in the MDVC in a locality served by a fixed vaccination point (cross in A), and by a mobile team (in B, X’s represent locations where the mobile team stopped to wait for dogs or to vaccinate dogs). Created with MapGAM package [38] in R [39].

## 4. Discussion

In Arequipa Peru, the social and spatial aspects of urbanization facilitate the emergence of dog rabies and complicate its control. In 21 urban and 9 peri-urban localities in Arequipa, Peru, we found low vaccination coverage and coverage that was spatially uneven. We found a strong effect of a potential proximal determinant of participation in the MDVC: distance to the vaccination point. The unadjusted data show a clear negative gradient with higher levels of participation in the MDVC at shorter distances to the vaccination point. After accounting for other important individual- and household-level variables, distance to the vaccination point remains an important factor associated with participating in the campaign. The association between distance to and participation in the MDVC also impacts the spatial evenness of vaccination. We found areas with statistically significant lower odds of dogs being vaccinated, and the LOESS smoother maps correlated well with maps of vaccination coverage. Therefore, it seems that both fixed point and mobile team canine vaccination approaches produced spatially heterogeneous vaccination coverage. However, we found that vaccination coverage was more “patchy” in localities served by mobile vaccination teams. This combination of mobile and fixed points was used also in 2015, but the same localities are not always served with the same approach (e.g. a locality that was served with mobile teams in 2015 could be served with fixed point vaccination in 2016). As others have reported [40], there is potential that in some localities owners in 2016 expected that the MDVC would be brought to their doors and did not plan or intend to bring the dogs to the fixed points in their areas. Spatially heterogeneous vaccination coverage is undesirable for dog rabies control and elimination. Townsend et al. [41] found that such patchy coverage can “profoundly damage prospects of elimination […] by creating pockets where rabies could persist”, and modeled patchy coverage within 1 km^2^ cell grids. The low coverage ‘patches’ in our study had smaller areas than 1 km^2^, thus the potential for a threat to elimination efforts may be different or non-existent. However, in these densely populated areas it is unknown if these ‘patches’ are large enough to sustain rabies transmission in the city.

Many studies have explored logistical, informational, social and structural barriers for dog rabies vaccination experienced by owners in rabies-affected areas [1,17,18,23,24,40,42–52]. Two of the most common reasons identified are difficulty handling the dogs [1,23,24,48,49,51,52] and lack of time [1,24,48,49,51,52]. These two logistical barriers are correlated with a less studied element: the distance to the vaccination campaigns. Distance to health services has been fairly well studied in terms of availability of and access to health care and impact on health, especially for maternal health, treatment and prevention of chronic diseases and treatment adherence for infectious diseases [53–55], but not as much for preventing infectious diseases. Some rural studies mention distance as a potential factor for low dog vaccination coverage [23,56] and two studies directly evaluated the association between distance and overall rural villages vaccination coverage [49] and the association between distance and attendance at the MDVC in Sub-Saharan Africa [18]. In the sub-Saharan Africa study, researchers found that distance in dispersed communities have an impact on MDVC attendance [18]. Given the hilly landscape with rare direct paths between houses and vaccination points, they estimated the shortest-path distance for their analysis to take into consideration the long and tortuous routes dog owners had to follow to visit the vaccination sites. In our study area, with non-dispersed urbanized highly-populated localities and high density of street intersections that increase walkability, distance is still an important proximal explanation for low participation in the vaccination campaigns.

Our study area consisted of urban and peri-urban localities. Surprisingly, there were no clear differences in participation in the MDVC between these two groups. However, the distribution of other proximal rabies-related characteristics is different between them (e.g. more free-roaming dogs in peri-urban areas, more neutered/spayed dogs in urban areas). There are other social determinants that provide more distal explanations for participation in the MDVC. In previous focus groups conducted by our team, young females reported that having a baby at home could prevent them from participating in the campaign [1]. Similarly, we found that households with children under 5 years old were less likely to vaccinate their dogs compared to houses without children, and each additional child under 5 reduced the odds of vaccinating the dogs in the house. This insight suggests an opportunity to increase participation in the MDCV by framing the decision to vaccinate as an action taken to protect children in the household from rabies. There was also a clear difference in participation between those who lived in a dog rabies-endemic area before living in Arequipa. A possible explanation for that difference is higher awareness among that group. In our focus groups, we found low awareness and low perception of severity among residents of Arequipa [1]. Interestingly, another component of migration history was also associated with participation in the MDVC: time living in the area. This phenomenon has been observed for the utilization of other health services in different settings and populations [53–55]. Migration, settlement and adaptation are processes that take time and are necessary for the uptake of health services [56–59], and could be influencing the participation in the MDVC. Importantly, in the peri-urban localities there are more households with children under 5, more recent migrants, and more people whose previous residence was in a rabies-affected region.

Our study has a number of limitations. Other studies have focused attention on dog-level variables (e.g. sex, age, function) that might be associated to participation in MDVC [18]. We did not analyze these; rather we focus on owner and community characteristics that can be utilized by the health authorities (who rarely have the opportunity to collect detailed house-by-house information) to increase participation in MDVC. Vaccination status and access to the street were reported by the owner/interviewee. Given the bad publicity in the local media about owned free roaming unvaccinated dogs and the authorities’ threats to fine ‘irresponsible’ dog owners, there is potential for social desirability bias to inflate our estimate of vaccination coverage and deflate that of the proportion of dogs that have access to the street. We did not ask the interviewees to show the vaccination certificate they receive at the MDVC since many of them do not save these certificates. We used Euclidean distances, which are only a proxy for the real distance traveled by individuals to a vaccination post.

Dog rabies virus transmission continues in Arequipa, putting at risk millions of people in the city and the surrounding departments. Dog-focused public health strategies are not limited to rabies: deworming dogs to prevent human echinococcosis [57], use of insecticide treatment [58] or vaccines [59] on dogs to prevent human Leishmaniasis or Chagas disease, are just a few examples. These programs, if they are to be successful, require high coverage and evenness in their implementation. The same approaches to reach the appropriate levels of community participation that might have worked in the 1980s are not working today. Understanding the barriers for dog owners’ participation in community-based programs will be crucial to implement effective zoonotic disease preventive activities. Distance to health services and the heterogeneous social composition of growing cities have to be examined when designing field programs to protect against zoonotic diseases.

## Supporting information

S1 Alternative Language Abstract and S1 Table

## 5. Acknowledgments

We thank all the survey participants for agreeing to be part of our study and sharing their time with us. We acknowledge the work of the members of the Zoonotic Disease Research Laboratory for their contribution to the implementation of this study, especially Katty Borrini and Jorge Apaza.

## 7. Supporting Information Legends

S1 Alternative Language Abstract. Resumen en español.

S1 Table. Reported exposure to MDVC communication channels by participation level.

