## Supplementary material for "Socio-spatial heterogeneity in participation in mass dog rabies vaccination campaigns, Arequipa, Peru": S1 Alternative Language Abstract and S1 Table

Supporting information

S1 Alternative Language Abstract

Para controlar y prevenir la rabia en Latinoamérica, se implementa campañas masivas de vacunación canina (CMVC) principalmente con puntos de vacunación fijos: los dueños deben llevar sus perros a los puntos de vacunación donde los perros son vacunados gratuitamente. La rabia canina todavía es endémica en algunos países de Latinoamérica y altas coberturas de vacunación y coberturas uniformes son atributos deseados en las CMVC para detener la transmisión del virus rábico. En Arequipa, Perú, realizamos encuestas puerta a puerta en más de 6000 casas luego de la CMVC (VANCAN en Perú) para evaluar la asociación de la ubicación de los puntos de vacunación y estos dos atributos. Encontramos que el chance de participar en la campaña disminuía en 16% por cada 100 metros entre la casa de los propietarios y el punto de vacunación más cercano (p=0.041), luego de controlar por potenciales confusores. Encontramos determinantes sociales asociados a la participación en la VANCAN: por cada niño menor de 5 años en la casa, el chance de participar en la VANCAN disminuyó en 13% (p=0.032), y por cada década menos de residencia en el área, el chance de participar en la VANCAN era 8% menor (p<0.001), luego de controlar por distancia y otras covariables. También encontramos agrupación espacial significativa de perros no vacunados, sobre los 500 m a los puntos de vacunación, que crearon “bolsones” de perros no vacunados que podrían mantener la transmisión de la rabia. Entender las barreras para la participación de los dueños de perros en programas comunitarios de vacunación canina será crucial para implementar actividades preventivas efectivas contra las enfermedades zoonóticas. Los elementos espaciales y sociales de urbanización tienen un rol importante en las coberturas de las CMVC y deberían ser considerados durante su planeamiento y evaluación.

S1 Table. MDVC communication channels interviewees were exposed by participation level.

| How they learned about the campaign | Vaccinate none dog (n=858) | Vaccinated some dogs (n=170) | Vaccinated all dogs (n=1,397) | p |
| --- | --- | --- | --- | --- |
| Megaphone | 56.9% | 64.5% | 66.5% | <0.001 ^a^ |
| TV | 36.4% | 33.1% | 32.1% | 0.842 ^a^ |
| Radio | 33.9% | 23.3% | 30.4% | 0.122 ^a^ |
| Poster/Banner | 7.8% | 9.9% | 7.4% | 0.337 ^a^ |
| Relative/friend/neighbor | 4.8% | 8.7% | 5.7% | 0.027 ^a^ |
| Newspaper | 1.3% | 0.6% | 1.1% | 0.798 ^a^ |
| Flier | 1.3% | 1.2% | 0.7% | 0.497 ^b^ |
| At municipality | 0.6% | 0.6% | 0.6% | -- |
| Social media | 0.1% | 0.6% | 0.7% | 0.066 ^b^ |
| Community meeting | 0.1% | 0.0% | 0.0% | 0.454 ^b^ |

p-values estimated with ^a^ Chi square test and ^b^ Fisher exact test.
